# RNase III cleavage signals spread across splice junctions enforce sequential processing of co-hosted snoRNAs

**DOI:** 10.1101/2025.01.08.631648

**Authors:** Valérie Migeot, Yves Mary, Etienne Fafard-Couture, Pierre Lombard, François Bachand, Michelle S Scott, Carlo Yague-Sanz

**Author notes:** Correspondence should be addressed to C.Y-S.

## Abstract

Small nucleolar RNAs (snoRNAs) are a class of non-coding RNA molecules whose precursor transcripts are capped and polyadenylated. However, these end modifications are detrimental to snoRNA function and must be removed, a process typically involving excision from introns and/or endonucleolytic cleavage. In the case of polycistronic RNA precursors that host multiple snoRNAs, the sequence of maturation events is not well understood. Here we report a new mode of maturation concerning snoRNA pairs that are co-hosted in the intron and the adjacent 3’ exon of a precursor transcript. For such a pair in the model eukaryotic species *Schizosaccharomyces pombe*, we identified a double-stranded RNA hairpin folding across the exon-exon junction. The hairpin recruits the RNase III Pac1 that cleaves and destabilizes the precursor transcript while participating in the maturation of the downstream exonic snoRNA, but only after splicing and release of the intronic snoRNA. We propose that such RNase III degradation signal hidden in an exon-exon junction evolved to enforce sequential snoRNA processing. Sequence analysis suggest that this mechanism is conserved in animals and plants.

## Introduction

Small nucleolar RNAs (snoRNAs) constitute a class of eukaryotic non-coding RNA that guide chemical modifications on other RNA molecules. Most snoRNA can be classified into one of two major snoRNA families based on distinct structural features, functions, and conserved sequence motifs (or boxes). The C/D box snoRNAs fold into a closed loop structure and associate with methyltransferase proteins to deposit 2′-O-methylation on target RNAs. In contrast, H/ACA box snoRNAs adopt a double hairpin structure and interact with pseudouridine synthase proteins to guide pseudouridination. Members of both snoRNA families specifically recognize their targets through base-pairing interactions between guide sequences in the snoRNA and complementary sequences in the substrate RNA^1,2^. One of the major targets for snoRNA-directed modification is ribosomal RNA (rRNA), which aligns with the preferential nucleolar localization of snoRNAs.

SnoRNA maturation and biogenesis in eukaryotes rely on complex mechanisms that are often perceived as unusual compared to other gene classes. Being transcribed by RNA polymerase II (RNAPII), snoRNA precursor transcripts (pre-snoRNAs) undergo co-transcriptional capping and polyadenylation, similarly to messenger RNAs (mRNAs). The 7-methylguanosine (m^7^G) cap and poly(A) tail are essential modifications that protect mRNA from exonucleolytic degradation and facilitate their export and translation in the cytoplasm^3^. In contrast, mature snoRNAs are intrinsically stable due to their structure and the association of partner proteins, and do not require end modifications –which can even be detrimental to snoRNA function. For instance, m^7^G-capped snoRNA in yeast have been shown mislocalize to the cytoplasm, where they are targeted for degradation^4^. Therefore, the m^7^G cap and poly(A) tail need to be removed or altered from pre-snoRNAs, relying on distinct mechanisms depending on snoRNA genomic organization: intronic, (mono)-exonic, or polycistronic, reviewed in^5^ and briefly summarized hereafter.

In mammals, snoRNAs are often encoded in introns of host genes. For instance, 61% of all annotated snoRNAs in human are intronic^6,7^. Following splicing, intronic snoRNAs can be excised from the lariat either by endonucleolytic cleavage or by the combined action of the debranching enzyme and exonucleases, resulting in cap- and poly(A)-less linear transcripts. Remarkably, intronic snoRNAs are often hosted in genes coding for proteins involved in ribosome synthesis, suggesting that snoRNA and ribosome production are coordinated. However, host genes are not exclusively protein coding: a significant fraction of intronic snoRNA (23% in human)^6,7^ are hosted in long non-coding RNAs (lncRNA) genes, most of which have no apparent function beyond serving as snoRNA precursors.

Exonic snoRNAs (*sensu lato*, referring to all snoRNA that are not intronic) are more common in yeasts, where they account for 90 and 81% of all snoRNAs in budding yeast and fission yeast, respectively^6^. They can be organized either as polycistrons or as individual genes. For individually encoded snoRNAs, 3’ end processing differ among species. In fission yeast, it is mediated by the mRNA cleavage and polyadenylation machinery^8^, with subsequent deadenylation by the nuclear exosome guided by the nuclear poly(A)-binding protein Pab2^9^. 5’-end processing differ between snoRNA families with most individual C/D box pre-snoRNAs being transcribed with a capped 5’ leading sequence which is subsequently removed by endonucleolytic cleavage and exonuclytic trimming^10,11^. Conversely, for pre-snoRNA transcribed without a leading sequence (including most H/ACA snoRNA in yeast), the m^7^G cap of the pre-snoRNA is converted into a trimethylguanosine (m^2,7,7^G or TMG) cap^12^.

Finally, exonic snoRNA genes can also be organized into operons, where snoRNAs are typically matured from polycistronic transcripts through endonucleolytic cleavage between the cistrons. In budding yeast, this process is mediated by the RNase III family protein Rnt1^10,13^. In other species, including fission yeast and human, the actors mediating the 5’ and 3’ maturation of exonic snoRNA co-hosted in the same precursor transcripts remain largely unknown. In these cases, the sequence of maturation events is particularly significant, as precursor cleavage may expose snoRNAs and surrounding sequences to exonucleolytic degradation. Whether and how the order of polycistronic snoRNA processing is controlled remains an open question.

In this study, we address these questions using the model eukaryote *Schizosaccharomyces pombe*. In *S. pombe*, the RNase III homolog Pac1 is an endoribonuclease involved in RNA polymerase II termination and ncRNA biogenesis, including the 5’ maturation of several exonic C/D box snoRNAs^14^. Mechanistically, Pac1 targets doublestranded RNA (dsRNA) hairpins with a minimum stem length of 20 base pairs, tolerating small bulges and varied loop structures^15,16^, and cleaves them in two staggered cuts. This contrasts with its homolog Rnt1 in budding yeast that specifically targets for cleavage RNA hairpins with apical NGNN tetraloops^17^.

Here, we report a new role for Pac1 in the maturation of a snoRNA pair co-hosted within the intron and adjacent 3’ exon of a precursor transcript. Remarkably, Pac1 activity at this locus depends on a dsRNA hairpin that folds across the exon-exon junction after splicing of the intron. Our findings suggest that this exon-exon junction serves as a conditional degradation signal that ensures sequential snoRNA processing, a mechanism that may be conserved across eukaryotes, including animals and plants.

## Results

### snoU14 and mamRNA are transcribed from a polycistronic precursor stabilized by Pac1 inactivation

Based on high-throughput genomic and transcriptomic data, we previously identified Pac1 cleavage signatures on at least five C/D box snoRNA in *S. pombe*, including snoU14, for which Pac1 inactivation lead to the accumulation of 5’-extended precursor transcripts^14^. Accordingly, we proposed that Pac1 is involved in snoRNA maturation through 5’-end processing^14^. However, the recent discovery by the Rougemaille group of a previously unannotated ncRNA named *mamRNA* (for *Mmi1 and Mei2-associated RNA*) directly upstream of snoU14^18^ suggests that Pac1’s role at this locus may be more complex than previously thought. Moreover, a recent preprint from the same research group describes that mamRNA and snoU14 are expressed from a common precursor transcript^19^, a conclusion we also reached when independently evaluating the contribution of Pac1 to mamRNA/snoU14 expression.

To assess Pac1’s role in modulating mamRNA expression, we first performed northern blot experiments using radiolabeled probes complementary to the mamRNA (*C*), snoU14 (*B*) and the intergenic region between the two genes (*A*) under conditions where the activity of Pac1 is compromised. Specifically, we used either the Pac1-AA (anchor-away) strain, which enable the conditional nuclear exclusion of Pac1, or the Pac1-ts (thermosensitive) strain, which carries a hypomorphic Pac1 allele that impairs Pac1 activity even at growth-permissive temperature^14^. As expected, the *A* and *B* probes recapitulated our previous observation that a long 5’-extended precursor containing the mature snoU14 sequence accumulates in conditions of Pac1 deficiency (Figure 1.A, *A*-*B* probes). More surprisingly, at least four distinct transcripts were detected with the mamRNA probe (Figure 1.A, *C* probe). Among these, the two shortest ones were the most abundant within the control strains (when Pac1 is active), with length matching the annotated mamRNA *short* and *long* isoforms^18,20^. In addition, a longer, previously uncharacterized mamRNA isoform specifically accumulated in the Pac1-deficient strains. Since this isoform migrates at the same size as the 5’-extended snoU14 precursor, we tentatively named it *pre-mamRNA/snoU14*. Long read RNA-sequencing validated the existence of such transcript isoforms containing both the mamRNA and snoU14 sequences (Supplementary Figure 1.A), as also recently reported^19^. Further suggesting that they are transcribed together, previously published Cap Analysis of Gene Expression (CAGE) data^21^ highlighted a single transcription start site for the entire locus, located directly upstream from the mamRNA gene (Figure 1.B). Collectively, these results confirm that snoU14 and mamRNA are transcribed from a single precursor^19^ and show that the abundance of this precursor depends on Pac1 activity.

**Figure 1:**
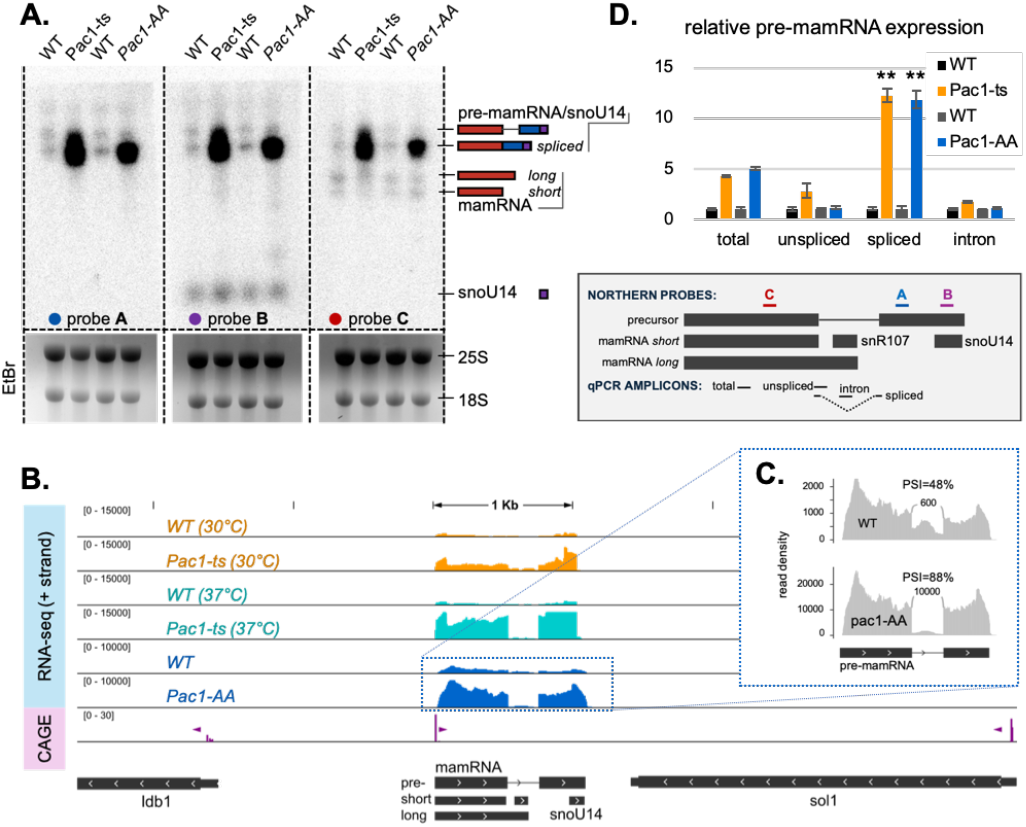
A common precursor for mamRNA and snoU14 depends on Pac1 activity. **A**. Representative northern blot analysis of transcripts originating from the mamRNA/snoU14 locus under conditions where the activity of Pac1 is compromised. Probes positions are indicated below panel D. **B**. Genome browser snapshot of the normalized read density over the mamRNA/snoU14 locus for RNA-seq experiments (+ strand only) and CAGE-seq (+ and - strands). For the CAGE-seq track, the direction of transcriptional firing is indicated with purple arrowheads. Gene annotation are shown in dark grey. **C**. Sashimi plot depicting the junctions reads as arcs over the read density profiles for the Pac1-AA and matching WT RNA-seq experiments at the mamRNA locus. The number of junction reads (rounded to the nearest 100) and percent spliced-in values (PSI) are indicated. **D**. RT-qPCR analysis (n=3) of transcript isoforms spanning the mamRNA locus. The positions of qPCR amplicons are indicated below the panel. Statistical significance of the differences in spliced isoform level in the Pac1 mutants compared to their respective control strains is indicated (**: Student’s *t*-test pvalue < 0.01). EtBr: Ethidium Bromide; ts: thermosensitive; AA: anchor-away; WT: wild-type.

We then corroborated our northern blot results with short read RNA-sequencing data, confirming that transcripts originating from the locus encompassing both the mamRNA and snoU14 genes were upregulated in Pac1-deficient strains (Figure 1.B). Notably, a 227-nucleotides gap in the read density profiles was observed within the mamRNA. This gap corresponds to a newly described intron in the pre-mamRNA/snoU14 transcript^19^ which is further confirmed here by spliced reads spanning the exon-exon junction in both short and long read RNA sequencing data (Figure 1.C, Sup. Figure 1.A). Interestingly, this intron hosts a newly annotated C/D box snoRNA, snR107^19^, but its expression remained largely unaffected by Pac1 inactivation (Sup. Figure 1.B).

Across the mamRNA intron, we observed that the spliced reads accumulated more than the intronic reads under Pac1-deficient conditions (Figure 1.B-C), suggesting that the isoform affected by Pac1 activity was spliced. In contrast, the steady-state level of the unspliced pre-mamRNA/snoU14 isoform (Figure 1.A) remained largely unchanged. To further confirm these observations, splice-specific RT-qPCR experiments showed a significant increase in relative expression for the amplicon spanning the exon-exon junction within the mamRNA, but not for the amplicons spanning either the exon-intron junction nor the intronic region (Figure 1.D). Therefore, we conclude that Pac1 activity specifically reduces the expression levels of the spliced pre-mamRNA/snoU14 isoform.

### A stemloop structure at the pre-mamRNA exon-exon junction directs Pac1 cleavage

We next wanted to identify *cis*-acting elements in the pre-mamRNA/snoU14 transcript responsible for Pac1 recruitment and/or activity. The sequence directly upstream of snoU14 has previously been predicted to fold in a stemloop which could serve as a Pac1 substrate; however, this has not been experimentally tested^14^. To challenge this prediction, we introduced eight point mutations expected to dramatically disrupt the predicted structure and named the resulting mutant *snoU14-SD* (*<u>S</u>tem<u>D</u>ead*) (Figure 2.A). In this strain, the mature snoU14 levels were significantly decreased by about 34% (Figure 2.A, *B* probe; quantified in Sup. Figure 2.A), suggesting that the mutated sequence might play a role in snoU14 expression. However, the snoU14-SD mutant did not phenocopy Pac1 deficiency: the expression of the spliced pre-mamRNA/snoU14 isoform remained at wild-type levels, as measured by both Northern blot and splice-specific RT-qPCR (Figure 2.B-C). From these observations, we conclude that the predicted stemloop proximal to snoU14 was not responsible for Pac1 activity on the precursor transcript.

**Figure 2:**
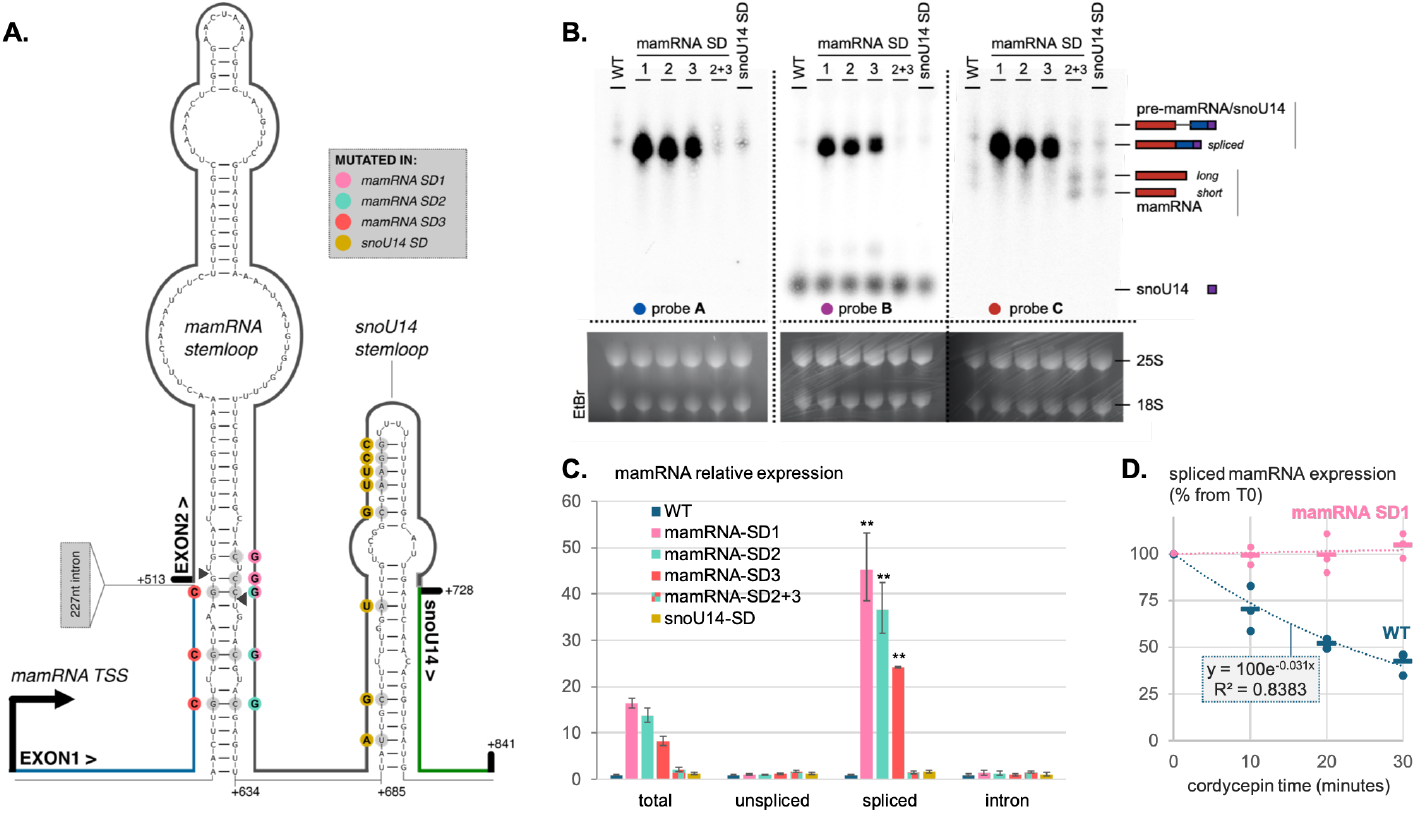
A stemloop spanning the exon-exon junction directs Pac1 cleavage at the mamRNA. **A**. Predicted secondary structures of the spliced pre-mamRNA/snoU14 transcript. Nucleotide positions are numbered relatively to the mamRNA transcription start site (TSS). Mutated bases are color-coded according to the StemDead (SD) mutants in which they were altered. Putative Pac1 cleavage sites, inferred from degradome-seq, are indicated by arrowheads. **B**. Northern blot analysis of transcripts originating from the mamRNA/snoU14 locus. Probes positions are the same as in Figure 1.A. EtBr: ethidium bromide. **C**. RT-qPCR analysis (n=3) of transcript isoforms spanning the mamRNA locus. The qPCR amplicons are the same as in figure 1.D. Statistical significance of differences in spliced isoform levels between CS mutants and their control strain (WT) is indicated (**: Student’s *t*-test pvalue < 0.01). **D**. Decay rate of the spliced pre-mamRNA/snoU14 transcript, measured by RT-qPCR after transcriptional inhibition with 0.6 mM cordycepin (n=3).

We then scanned both the unspliced and spliced pre-mamRNA/snoU14 sequences for other RNA structures that could potentially serve as Pac1 substrate. Strikingly, a very stable stemloop (ΔG=-35Kcal/mol) with symmetric bulges was predicted to fold across the exon-exon junction (-15 of exon 1 to +121 of exon 2) of the spliced precursor (Figure 2.A). Suggesting that this stemloop could constitute a genuine Pac1 substrate, degradome-seq experiments^22^ revealed that the 5’-ends of two cleavage intermediates (Sup. Figure 2.A) map on its structure and show the atypical 2nt-overhand signature of RNase III cleavage (Figure 2.A, arrowheads). Importantly, this stemloop was not predicted in the unspliced isoform, which instead formed a branched structure due to the disruptive presence of the intron (Sup. Figure 2.A).

To verify that the identified stemloop was responsible for Pac1 activity at the locus, we introduced three sets of point mutations in the endogenous mamRNA gene, generating the *mamRNA-SD*_*1*_, *-SD*_*2*_, and *-SD*_*3*_ mutants (Figure 2.A). The disruptive mutations of the *mamRNA-*SD_1_ and SD_2_ mutant are located in the exon 2, near the 3’ end of the predicted structure. Conversely, the disruptive mutations in the mamRNA-SD_3_ mutant are located in the exon 1, near the 5’ end of the predicted structure. The SD_2_ and SD_3_ mutations were designed so that their combination restores base-pairing, reforming the structure that the individual mutations disrupted, and therefore allowing us to discriminate between sequence- and structure-dependent effects.

In all individual mamRNA-SD mutants (mamRNA-SD_1_, SD _2_, and SD_3_), the steady-state levels of the spliced pre-mamRNA/snoU14 isoform were dramatically increased (Figure 2.B-C) while the mature mamRNA forms were less abundant (Figure 2.B, *C* probe ). Strikingly, these effects were not observed in the mamRNA-SD_2+3_ mutant where the SD_2_ and SD_3_ complementary mutations reform a stable structure. These results strongly suggest that the RNA secondary structure spanning the exon-exon junction of mamRNA is responsible for the Pac1-mediated repression of the spliced pre-mamRNA/snoU14 isoform. Confirming this hypothesis, we found that mutations disrupting the secondary structure were epistatic to Pac1 loss of function, as Pac1 exclusion from the nucleus in the mamRNA-SD1 background did not further increased the levels of the spliced pre-mamRNA/snoU14 isoform (Sup. Figure 2.A).

Arguably, the most straightforward explanation for how Pac1 represses the *pre-mamRNA/snoU14* isoform is that its endonucleolytic activity destabilizes the transcript. To test this hypothesis, we measured the decay rate of mamRNA isoforms in a time-course experiment following transcription inhibition with the chain-terminating adenosine analog cordycepin (see material and methods section for details about inhibitor selection). In wild-type cells, cordycepin treatment caused a rapid decrease in the relative abundance of the spliced mamRNA/snoU14 precursor (estimated half-life assuming a complete transcriptional inhibition = 20 minutes). However, in the mamRNA-SD_1_ mutant, the isoform was drastically stabilized, with no detectable decay over the course of the experiment (Figure 2.D). From these results, we concluded that the stemloop structure over the splice junction destabilizes the spliced mamRNA-snoU14 precursor in a Pac1-dependent manner.

### snoU14 can mature independently from Pac1 activity

The degradation signal localized at the mamRNA splice junction raises the question of its role in snoRNA maturation. We initially hypothesized that Pac1 cleavage at this locus was necessary to release snoU14 from the mamRNA precursor. Supporting this idea, Northern blot experiments showed that snoU14 is sequestered in the longer precursor transcript when Pac1 activity is compromised in Pac1-deficient strains or mamRNA SD mutants (Figure 1A and 2B, *B* probe). However, mature snoU14 levels remained unaffected, suggesting that alternative pathways are able to mature snoU14 from its precursor, potentially with slower kinetics.

SnoU14 is a conserved and essential C/D box snoRNA required for the maturation of the 18S rRNA from the polycistronic 35S rRNA precursor. Accordingly, deficiencies in snoU14 expression have been reported to severely affect rRNA processing, 18S levels, and cellular growth in yeast^23,24^. Here, northern blot analysis on rRNA precursors showed that reduced snoU14 expression in the snoU14-SD strain (Figure 2B) correlated with a modest (∼20%) but statistically significant increase in 35S and 33/32S rRNA precursor (Sup. Figures 3.A-B). In contrast, no perceptible differences in mature or precursor 18S rRNA levels could be observed in the mamRNA-SD mutants (Sup. Figure 3.A-B), indicating that snoU14 matured by putative Pac1-independent pathway remain fully functional. Consistently, all the mamRNA SD strains grew normally in rich medium (Sup. Figure 3.A). From these experiments, we conclude that Pac1 cleavage on the pre-mamRNA/snoU14 transcript is not strictly required for snoU14 expression or function.

**Figure 3:**
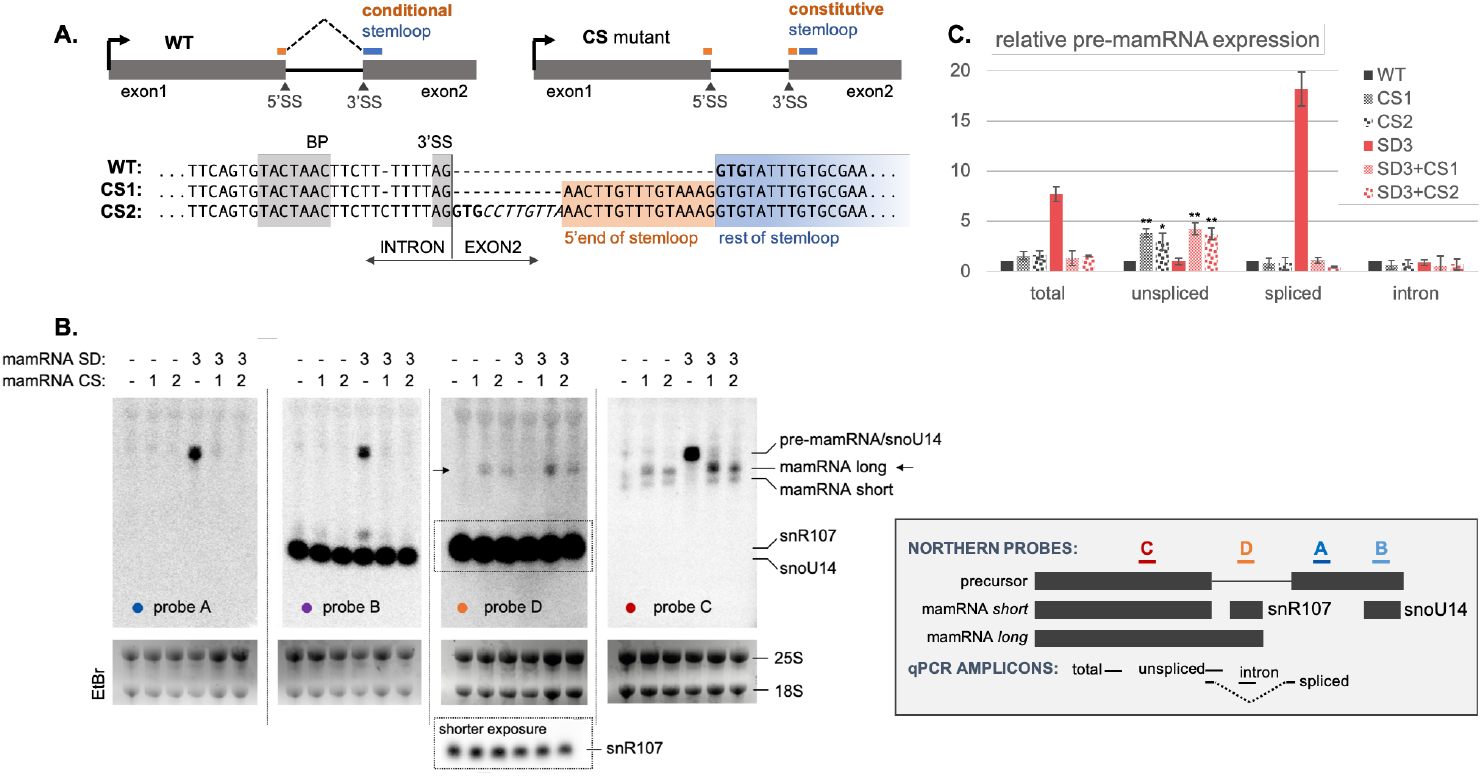
Conditional Pac1 cleavage ensures sequential snoRNA processing. **A**. Top: schematic representation of the mamRNA Constitutive Stem (CS) mutations. Bottom: mutated sequences of the mamRNA CS mutants. For the mamRNA-CS_2_ mutant, the nucleotides in bold and italic correspond to the original WT sequence immediately downstream the 3’ splice-site and immediately upstream the 5’end of the stemloop, respectively. SS: splice-site; BP: branch point **B**. Northern blot analysis of transcripts originating from the mamRNA/snoU14 locus. Probes positions are indicated on the right. Arrows indicates the snR107-containing long mamRNA isoform. EtBr: ethidium bromide. **C**. RT-qPCR analysis (n=3) of transcript isoforms spanning the mamRNA locus in the indicated strains. Statistical significance of the differences in unspliced isoform level in the CS mutants compared to their control strains is indicated (*: Student’s *t-*test p < 0.05; **: p < 0.01).

### Pac1 cleavage across splice junction coordinates the sequence of snoRNA maturation events

Disjoint secondary RNA structure spanning splice junctions are not unique to *S. pombe*; they can also be predicted upstream of snoU14 genes of related *Schizosaccharomyces* species, suggesting evolutionary conservation (Sup. Figure 4.A-B). We reasoned that such peculiar arrangement forces RNase III cleavage to occur only after intron splicing, a condition that may be required for proper expression of the cohosted intronic snoRNA. To test this hypothesis in *S. pombe*, we re-engineered the mamRNA/snoU14 locus by inserting a copy of the last 15 nucleotides of the exon1, corresponding to the 5’ end of the stemloop, at the beginning of the exon2 (figure 3.A). In the resulting strain, mamRNA-CS_1_ (*Constitutive Stemloop 1*), folding of the hairpin should be uncoupled from intron splicing, potentially leading to faster or unconditional Pac1 cleavage. An alternative version of the mutant (mamRNA-CS_2_), preserves the original sequence immediately downstream of the 3’ splice-site and immediately upstream of the 5’end of the hairpin (Figure 3.A). Strains carrying either the mamRNA-CS_1_ or -CS_2_ mutations grew normally in rich medium (Sup. Figure 3.A). Importantly, these mutations suppressed the increased levels of spliced precursor in the SD_3_ mutant (Figure 3.B, *A*-*B* probes; Figure 3.C), demonstrating that the re-engineered “constitutive” stemloops act as functional cleavage substrates for Pac1.

**Figure 4:**
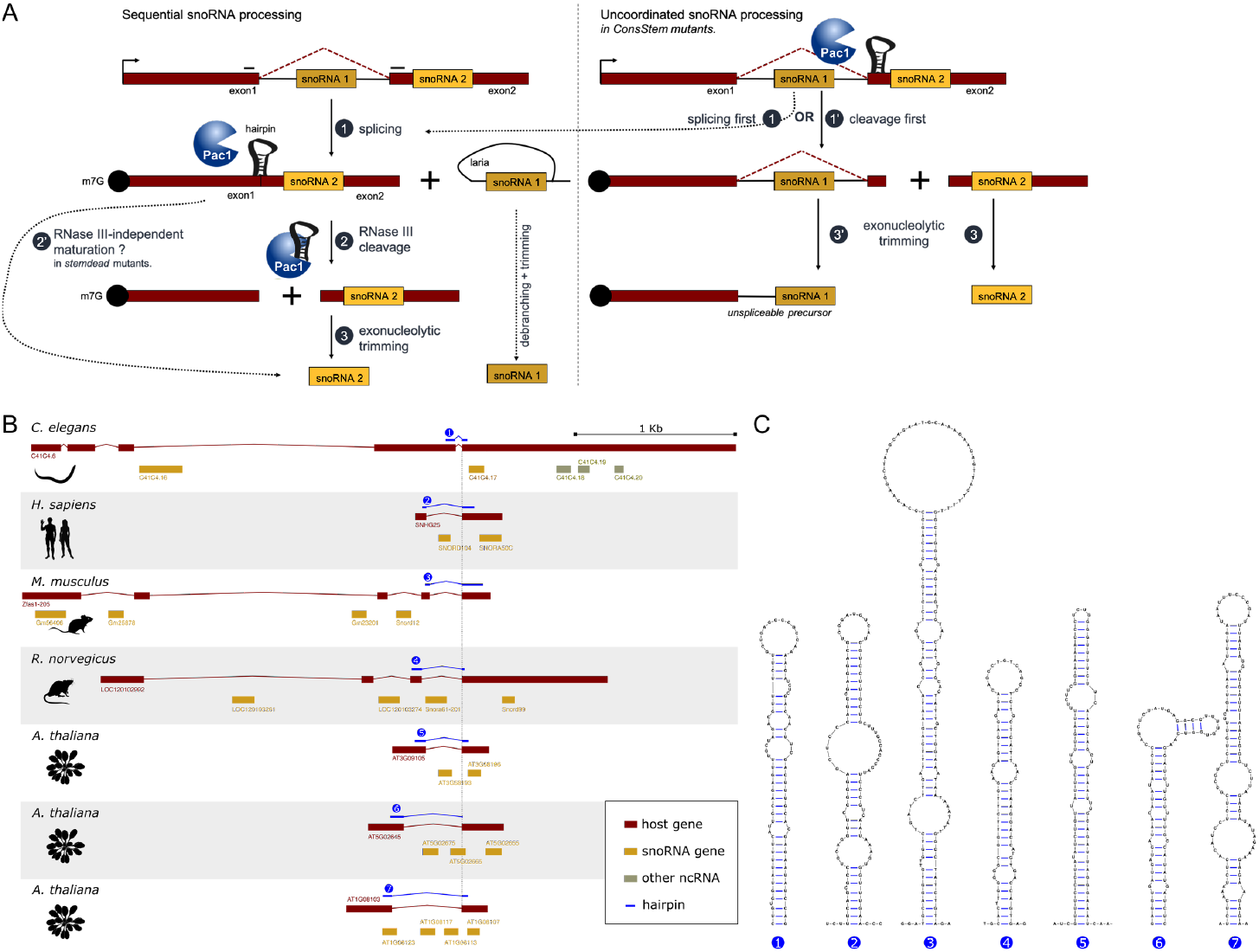
Sequential snoRNA processing of co-hosted intronic-exonic snoRNAs. **A**. Model for sequential vs uncoordinated processing of cohosted snoRNAs. In those models, m^7^G cap (black circles), polyA tails, and snoRNAs are barrier to exonucleolytic trimming. In sequential processing (left panel), splicing occurs first (1) allowing the release and maturation of the intronic snoRNA. By bringing together the two exons, splicing also allows for a secondary structure to fold across the exon-exon junction. This structure recruits the RNase III Pac1 in fission yeast (2), leading to the cleavage of the spliced precursor and subsequent maturation of the exonic snoRNA by exonucleolytic trimming (3). In the case of snoU14 in *S. pombe*, a secondary Pac1-independent maturation pathway exists (2’). In contrast, snoRNA processing would become uncoordinated (right panel) with cleavage potentially occurring before splicing (1’) if the RNA secondary structure folded constitutively. This could lead to the exonucleolytic degradation of the 3’ splice-site, leaving the intronic snoRNA trapped in the precursor (3’). **B**. Genomic arrangements of selected genes host to both intronic and exonic snoRNAs, aligned to the 5’ end of their last exon. **C**. Predicted secondary structure from the sequences spanning the last exonexon junction of host genes displayed in panel B.

Northern blot analysis of the CS mutants revealed that a small fraction of intronic snR107 remained trapped in a longer RNA species (Figure 3.A, *D* probe) whose 3’ end terminates two nucleotides downstream of snR107, as shown by 3’ RACE (Sup. Figure 3.A). This transcript therefore ressembles the previously annotated mamRNA *long* isoform^18,20^, though it is 15 nucleotides shorter, and contributes to the apparent increase in its expression in CS mutants (Figure 3B, *C* probes). These findings suggest that unconditional Pac1 cleavage can cause aberrant snR107 processing, although not to an extent sufficient to reduce mature snR107 level. A possible explanation for this observation is that Pac1 cleavage stochastically occuring before splicing on the mamRNA/snoU14 precursor could lead to 3’->5’ exonucleolytic trimming of the 3’ splice-site, antagonizing splicing and subsequent maturation of snR107 (Figure 4.A). Consistent with this model, qPCR analysis showed significantly increased levels of unspliced isoforms in the mamRNA CS strains (Figure 3.A).

### Conservation of secondary structures spanning splice junctions in mixed snoRNA clusters

Our observations suggest that a cleavage signal spread across splice junctions enforces sequential processing of intronic-exonic snoRNA pairs co-hosted within the same precursor transcript (Figure 4.A). We next wondered whether a similar mechanism might also exist in other eukaryotes. To address this question, we mined the *Ensembl* genome annotation^25^ for similarly organized snoRNA pairs, focusing on seven model species known for their high quality annotation: *S. cerevisiae, Caenorhabditis elegans, Drosophila melanogaster, Mus musculus, Rattus norvegicus, Arabidopsis thaliana*, and *Homo sapiens*.

In total, we identified twelve snoRNA clusters co-hosted within the same precursor transcript, each containing at least one exonic snoRNA and one intronic snoRNA. No such clusters were present in the intron-poor *S. cerevisiae* genome, whereas between one and three were found in the other species considered (Supplementary table 1). These findings suggest that, although relatively rare, *mixed* (exonic-intronic) snoRNA hosts exist in species beyond *S. pombe*. Among the identified clusters, the majority (ten out of twelve) are organized such that the last hosted snoRNA is exonic, preceded by one or more intronic snoRNAs (Figure 4.A). Of these ten clusters, at least seven harbored predictable RNA stemloop structures folding from the sequences spanning the last two exons of their spliced host precursor (Figure 4.C). Further work will be needed to determine whether these secondary structures also function as degradation signals and coordinate the maturation of co-hosted snoRNA.

**Table 1:** Consequences of RNase III cleavage site localization relative to intron boundaries.

| Localization | Consequences of cleavage | Shown in |
| --- | --- | --- |
| Within intron | quality control vs mis-spliced transcripts, regulation of targeted mRNA expression | *S. cerevisiae* (Rnt1) |
| Exon-intron junction | competition with splicing, antagonizes splicing | mammals (Drosha) |
| Exon-exon junction | splicing as a prerequisite to cleavage, delayed cleavage | *S. pombe* (Pac1) mammals ? |

## Discussion & Conclusions

In this study, we have demonstrated that in *S. pombe*, snoU14 and its upstream non-coding RNA, the mamRNA^18^, are expressed from a common polycistronic precursor whose transcription initiates approximately 1kb upstream of snoU14. The updated transcript model independantly confirm the findings of a recently posted preprint^19^ and provides a straightforward explanation for earlier observations that *S. pombe* snoU14 is functional when expressed from a *S. cerevisiae* promoter but cannot be expressed in *S. cerevisiae* in its native context, *i. e*. preceded by 800 bp upstream^23^. To explain this discrepancy, Samarsky and colleagues hypothesized that specific transcription signals from *S. pombe* snoU14 were poorly utilized in *S. cerevisiae*. However, it now appears that promoter elements, located further upstream, were simply not included in the transgene sequence.

Multiple transcripts arise from the mamRNA-snoU14 precursor, including an exonic snoRNA (snoU14), an intronic snoRNA (snR107)^19^, and both long and short mamRNA isoforms^18^. This diversity suggests that multiple RNA maturation processes coexist at the locus. Here, we demonstrated that the endoribonuclease Pac1 participates in the release of mature snoU14 from the precursor (Figure 1) However, this activity is dispensable for snoU14 maturation (Figure S3), hinting at a secondary maturation pathway which likely involves another endoribonuclease. Similarly, in *S. cerevisiae* where snoU14 is released from a dicistronic precursor through cleavage by the RNase III ortholog Rnt1, partial functional redundancy between Rnt1 cleavage and another unknown factor has been reported^26^. This raises the possi ,bility that the homolog of this unknown factor also operates in *S. pombe* as part of a secondary snoU14 maturation pathway.

We identified that Pac1 cleaves the mamRNA-snoU14 precursor at a stemloop RNA structure encoded disjointly over its two exons (Figure 2). The particular localization of this degradation signal is remarkable, as it makes Pac1 activity specific towards the spliced version of the precursor transcript, delaying cleavage until splicing is complete. This unique mechanism explains the dramatic accumulation of mamRNA-snoU14 precursor previously reported in splicing-deficient mutants^19^. We propose that requiring splicing as a prerequisite for Pac1 cleavage enable the precise control over the sequence of processing events. Indeed, we found that unconditional Pac1 cleavage of the mamRNA-snoU14 precursor leads to the accumulation of snR107-containing unspliced transcripts (Figure 3), suggesting that premature cleavage occurring before splicing can antagonize the later— possibly by exposing the 3’ splice site to degradation (Figure 4A). It is likely that core snoRNP proteins bound to snR107 sequence^19^ can shield what remains of the precursor from further exonucleolytic trimming, similar to the lncRNA transcripts with snoRNA ends (sno-lncRNAs) described in mammals^27^ and to hybrid messenger RNA–snoRNA (hmsnoRNA) described in *S. cerevisiae*^28^. Therefore, we propose that enforcing delayed cleavage for exonic snoU14 maturation ensures proper intronic snR107 maturation, which is critical to avoid the accumulation of a snR107-containing mamRNA isoform. Control over the expression of this isoform might be especially important, as both the mamRNA and snR107 contribute to the regulation of gametogenesis in fission yeast by interacting with the Mmi1 and Mei2 proteins^18,19^.

Previous studies have also described close mechanistic relationships between eukaryotic RNase III activity and intron splicing. In *S. cerevisiae*, Rnt1 cleavage sites are located within introns of seven pre-mRNAs coding for RNA binding proteins, promoting the degradation of unspliced pre-mRNA and lariat introns^29,30^. In mammals, Drosha cleavage sites span exon-intron junctions, antagonizing splicing either by direct cleavage or by sterically hindering splice-site recognition by the splicing machinery^31–34^. Our findings introduce a new, distinct configuration, where RNase III cleavage site spans exon-exon junctions and requires prior splicing (Table 1). A similar case has been briefly reported among the non-canonical Drosha substrates identified by genome-wide fCLIP-seq (formaldehyde cross-linking and immunoprecipitation) in human cell lines^35^.

The enforced sequential snoRNA processing mechanism described in this study may have evolved to accommodate the unusual genomic organization of the two co-hosted snoRNAs from the mamRNA locus, where one snoRNA (snR107, intronic) depends on splicing for its biogenesis^19^, whereas the other (snoU14, exonic) relies on endonucleolytic cleavage. Such an organization, which we classify as *polycistronic-mixed*, can be found in a wide range of eukaryotic species, though it is relatively uncommon compared to other types of snoRNA clusters^6^. However, its prevalence may be underestimated, as detecting polycistronic-mixed snoRNA clusters depends on highquality snoRNA and host RNA annotation. Ongoing efforts to improve these annotations using machine learning and RNA-sequencing optimized for snoRNA detection (Fafard-Couture et al., manuscript in preparation) may enhance classification accuracy for snoRNA organization.

Nevertheless, the polycistronic-mixed snoRNA clusters identified in the current annotation were preferrentially organized as a single exonic snoRNA located downstream of one or multiple intronic snoRNAs (Figure 4B). This consistent genomic organization, together with the presence of predictable stemloop structures folding across the last splice junction (Figure 4C), suggests that a cleavage signal spanning two exons may be a conserved mechanism to delay cleavage-dependent exonic snoRNA maturation until after the last intronic snoRNA of the cluster has been spliced out. Since the mechanism of 5’-end processing for exonic snoRNAs are unknown in higher eukaryotes, it remains to be determined which endonuclease(s) are involved (if any) and whether the predicted stemloops in the polycistronic-mixed clusters can indeed function as cleavage signals. Thus, the mechanisms underlying 5’-end processing of exonic snoRNAs in eukaryotes remain a promising area for future investigation.

## Acknowledgements

We thank Damien Hermand, Olivier De Backer, Marc Hennequart, Mathieu Rougemaille, and François-Xavier Stubbe for their helpful discussions and/or for proofreading the manuscript. We further thank Mathieu Rougemaille for generously sharing unpublished data regarding the existence of intronic snR107 in the mamRNA precursor. We also thank Damien Coupeau for his technical advice in setting up the 3’ RACE experiments. C.Y.-S. is a scientific collaborator of the Fonds de la Recherche Scientifique – FNRS.

## Author contributions

C.Y-S designed the study and wrote the manuscript with inputs from M.S and F.B. V.M, Y.M, and C.Y-S performed all experiments, with the exception of the nanopore sequencing experiment performed by P.L. E.F-C and C.Y-S analyzed transcriptomic and genomic data. C.Y-S prepared the figures. All authors reviewed the first draft of the manuscript.

## Materials and Methods

### Yeast strains, drugs and media

Unless stated otherwise, yeast cells were grown at 32°C in liquid minimal medium supplemented with adenine, uracil, histidine and leucin until an optical density of 0.4-0.6. For the anchor-away experiments, yeast cultures were treated for two hours with 2.5 µg/ml rapamycin (Sigma R8781) or equal volume of DMSO carrier to allow for Pac1 nuclear exclusion. For the estimation of RNA decay rates, yeast were grown at 26°C in EMM+AS over the full course of the experiment. Cordycepin (Sigma C3394, 10 mM stock in H_2_O, required heating at 55°C to fully dissolve) was added to a final concentration of 0.6 mM at the appropriate time so that all sample from the same experiments could be pelleted together.

Gene disruptions and C-terminal tagging of proteins were performed by PCR-mediated gene targeting using the lithium acetate method. Cloning-free CRISPR/Cas9-mediated mutagenesis was used to generate the stemloop mutants^36^. Yeast strains used in this study are listed in Supplementary Table 2.

### About the choice of cordycepin as a transcriptional inhibitor

Chemical inhibition of transcription in yeast is most commonly performed by using divalent cation chelator such as 1-10 phenanthroline or thiolutin^37,38^. However, divalent cation are required for other processes beyond transcriptional elongation by the RNA polymerases; they are also important for the spliceosome to catalyze the second step of splicing^39^ as well as for Pac1 cleavage activity^40^. Therefore, these transcriptional inhibitors interfere at different levels with the expression of the pre-mamRNA/snoU14, making them ill-suited to study its decay rate. As an alternative, we used the adenosine analog cordycepin. Compared to adenosine, cordycepin lacks an hydroxyl group required for chain elongation and therefore blocks transcription elongation when its triphosphorylated form is incorporated into nascent RNA^41^.

### RNA extraction and analysis

Total RNA was extracted following the classical hot phenol method^42^ as previously described^43^. Briefly, the cells were first resuspended in 750 µl of TES solution (10mM Tris-HCl [pH 7.5], 10mM EDTA [pH8.0] and 0.5% SDS) and 750 of acidic phenol-chloroform-isoamyl alcohol 125:24:1 (Sigma P1944) then incubated at 65 °C for 60 minutes with high-speed agitation for 10 seconds every 10 minutes. After centrifugation, the upper, aqueous, phase was transferred to a new tube containing equal volume of acidic phenol-chloroform-isoamyl alcohol 125:24:1. After thorough mixing and centrifugation, the upper phase was transferred to a new tube containing equal volume of chloroform-isoamyl alcohol 24:1 (Sigma 25666). After mixing and centrifugation, 500 µL of the upper phase was precipitated with 1.5 mL 100% ethanol and 50 µL of 3M NaOAc pH5.2. The precipitated RNA pellet was washed twice with 70% ethanol, air-dried and resuspended in DEPC-treated (Sigma D5758) water.

For RT-qPCR analysis, 0.5μg of total RNA was retro-transcribed with the High-Capacity cDNA Reverse Transcription Kit (Thermo 4368813) following the manufacturer’s instructions. The real-time PCR amplification was performed with SYBR Green 2X Supermix (Bio-Rad 172-5124) in a Bio-Rad CFX96™ Real-Time machine. The PCR program was 3 minutes at 95 °C followed by 40 cycles of (15’’ at 95 °C; 30’’ at 60 °C) and a melt curve. The PCR reactions were performed in technical duplicates, and all experiments were performed at least in biological triplicates.

Relative RNA quantification relied on the 2-^ΔΔCq^ method using the actin gene as internal reference except for the transcription inhibition assays (with cordycepin) in which we used the stable 18S rRNA instead. For statistical analysis, the normalized relative expression levels were brought back in the Cq scale by log2 transformation and used in one-sample Student’s *t*-test. Primer sequences are listed in Supplementary Table 3.

For northern blot analysis, two types of denaturing gel were used: 1.5% agarose gel in MOPS buffer with 1.6M formaldehyde to separate (pre-)snoRNAs, as previously described^18^, and 0.8% agarose gel in Tri/Tri buffer with 0.4M formaldehyde^44^ to separate pre-rRNAs, as described^45^. After migration, RNAs were transferred onto a positively charged nylon membrane (Amersham Hybond N+, Cytiva), crosslinked with UV in a Stratalinker or with heat (2h in the oven at 80°C), and pre-hybridized in hybridization buffer (Sigma H7033) at 42 °C. DNA probes were 5’ radiolabeled with [γ-32P]-ATP using T4 polynucleotide kinase (Thermo Scientific EK0031, 30 minutes at 37°C) and were added to the membrane. After over-night incubation, membranes were washed once with 2×SSC/0.1% SDS and twice with 0.1×SSC/0.1% SDS. A storage phosphor screen was exposed to the membrane for up to 24h and scanned in a Typhoon biomolecular imager (Amersham). When relevant, the samples were run in duplicates or triplicates on the same gel to allow the study of multiple probes without stripping. All northern blot experiments were performed at least in duplicates, with only one representative scan shown. Probe sequences are listed in Supplementary Table 3.

For 3’ RACE analysis, 10 µg of total RNA was polyadenylated using 5 U of *E coli* polyA polymerase (NEB) during 20 minutes at 37°C in presence of RNase inhibitor. After phenol-chloroform purification, 1 µg of polyadenylated and non-polyadenylated total RNA were reversed-transcribed with SuperScript III reverse transcriptase (Thermo Fisher Scientific) from an oligo-dT primer according to manufacturer’s instruction. RNA was degraded with RNase H and the cDNA was used in PCR reactions with a gene specific forward primer, a universal reverse primer and TAQ polymerase (with 45 seconds elongation and an annealing temperature of 61°C ). PCR products were ligated into pGEM-T easy vectors and electroporated in competent *E. coli* cells. Inserts were verified by PCR and sequenced.

For long read sequencing direct RNA-seq polyA+ libraries (SQK-RNA002, Oxford Nanopore Technologies) were prepared according to manufacturer’s instructions starting from 500ng of total RNA. The libraries were sequenced on a MinION apparatus during 48 hours.

### Sequence analysis

RNA secondary structures were predicted using the RNAfold2 webserver^46^ and visualized in vaRNA^47^. Short read RNA-seq data and degradome-seq data were processed as previously described^14,22^. For long read RNA-seq, the raw signal was basecalled with the Guppy algorithm provided by ONT (version 6.5.7) with options `--flowcell FLO-MIN106 --kit SQK-RNA002 --device auto --calib_detect --reverse_sequence --u_substitution`. The resulting reads were mapped onto S. pombe genome using minimap2^48^, version 2.24, with options -Y -G 1000 -t 8 -R “@RG\tID:Sample\tSM:hs\tLB:ga\tPL:ONT” --MD - ax splice -uf -k14 --junc-bed and visualized in the IGV software^49^.

## Data availability

Short read RNA-seq data and degradome-seq data re-analyzed in this study are available on GEO under the accession numbers GSE167041 and SRR12004691, respectively. Long read RNA-seq data analyzed in this study are available on ENA under the accession number PRJEB83877.

**Supplementary Figure 1:**
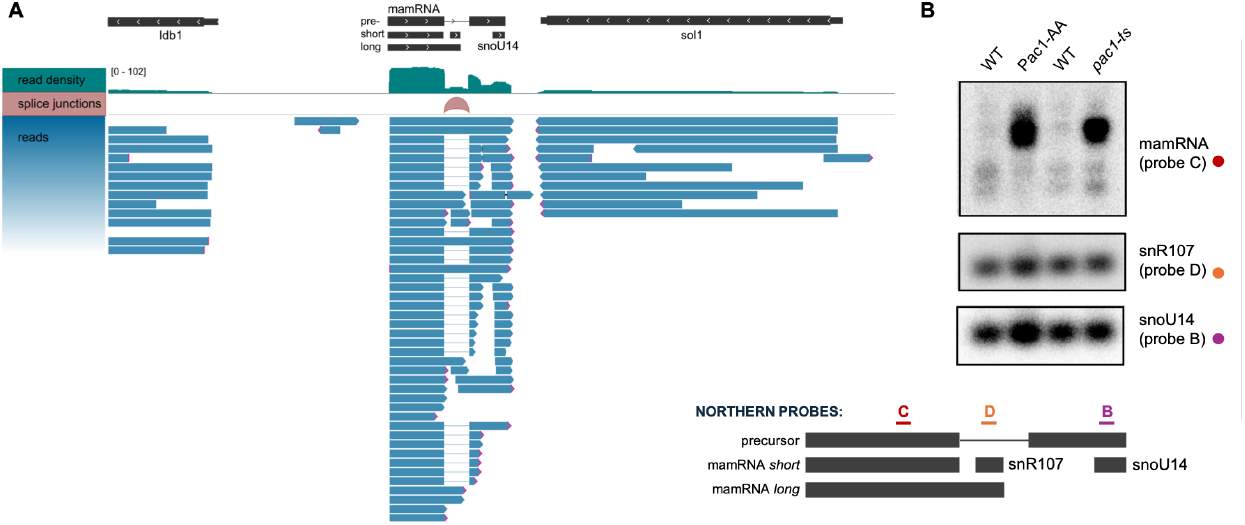
**A**. Read density, splice-junctions and individual reads from a long read sequencing experiment (wild-type strain) over the mamRNA/snoU14 locus. **B**. Representative northern blot analysis of transcripts originating from the mamRNA/snoU14 locus in conditions where the activity of Pac1 is compromised. Probes positions are indicated below panel B.

**Supplementary Figure 2:**
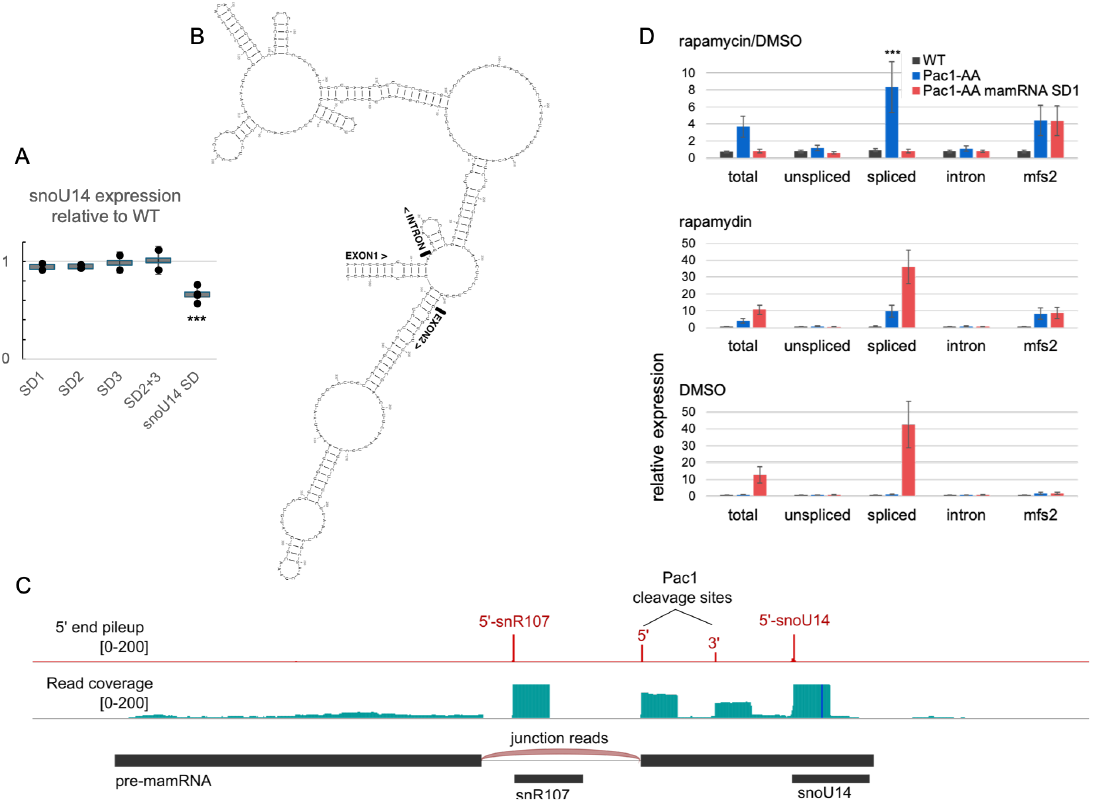
**A**. Semi-quantitative determination of mature snoU14 relative expression from pixel measurements of the northern blot experiments presented in Figure 2.A. Individual replicates are represented by filled circles. Means +/-SD are depicted by grey rectangle with error bars. Statistical significance of the differences in mature snoU14 level in the snoU14-SD mutant compared to the WT strains is indicated (***: Student’s *t*-test pvalue < 0.001). **B**. Predicted secondary structures of the unspliced pre-mamRNA/snoU14 transcript. **C**. 5’-end pile-up of reads and overall coverage from a degradome-seq experiment in wild-type strain (SRR12004691^22^) over the mamRNA-snoU14 locus. The putative 5’ and 3’ Pac1 cleavage sites are indicated. **D**. RT-qPCR analysis (n=3) of transcript isoforms spanning the mamRNA locus in the indicated strains treated for two hours with rapamycin (middle panel) or with its solvent (DMSO) as control (bottom panel). The qPCR amplicons are the same as in figure 1.A with the addition of mfs2 as a positive control – mfs2 have been previously shown to be upregulated by conditional nuclear exclusion of Pac1 by rapamycin in the Pac1-AA strain (Yague-Sanz et al, 2021). The fold change and statistical significance of the rapamycin treatment effect compared to the DMSO control are indicated for the spliced pre-mamRNA isoform (top panel, ***: Student’s *t*-test pvalue < 0.001).

**Supplementary Figure 3:**
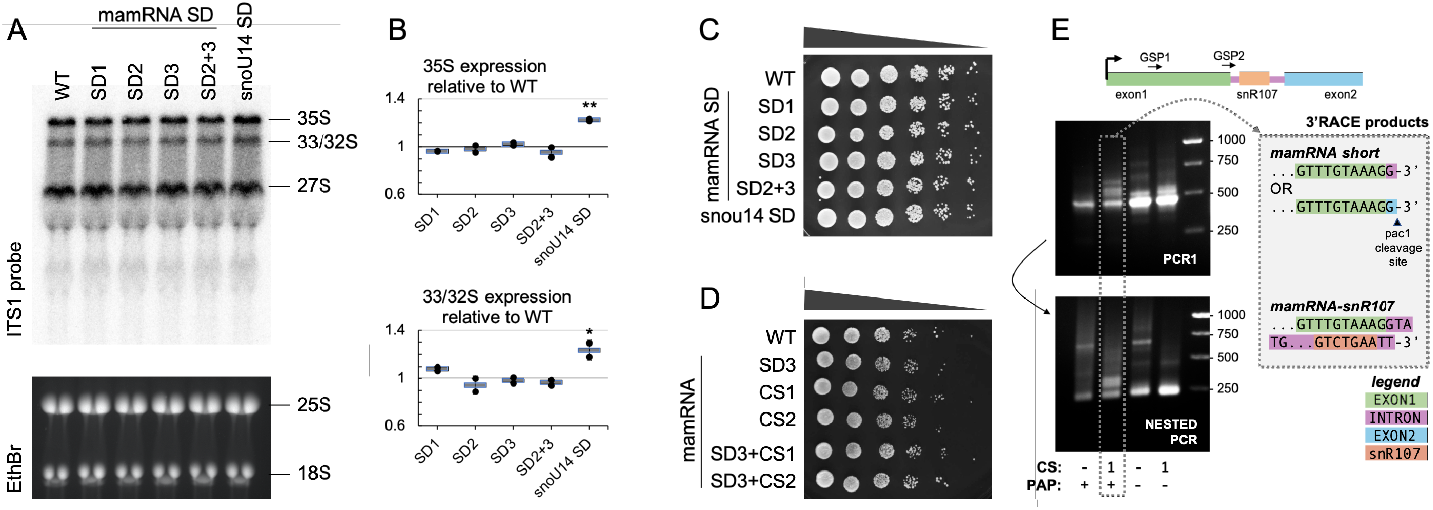
snoU14 matured independently from Pac1 is functional. **A**. Northern blot analysis of pre-rRNA transcripts in the indicated strains with internal transcribed spacer (ITS) probe. Mature rRNA forms (25S and 18S) are revealed by ethidium bromide (EthBr) staining. **B**. Semi-quantitative determination of mature 35 and 33/32S rRNA precursor relative expression from pixel measurement of the northern blot experiments presented in panel A. Individual replicates are represented by filled circles (n=2). Their mean +/-their standard deviation are depicted by grey rectangle with error bars. Statistical significance of the differences in snoU14-SD mutant compared to the WT strain is indicated (*: Student’s *t*-test pvalue < 0.05; **: Student’s *t*-test pvalue < 0.01). **C**. 5-fold dilutions of yeast cultures from the indicated strains spotted on YES-agar plates and incubated for 2 days at 32°C. **D**. 10-fold dilutions of yeast cultures from the indicated strains spotted on YES-agar plates and incubated for 3 days at 32°C. **E**. 3’-RACE analysis of mamRNA isoforms from total RNA extracted from CS1 mutant (CS) treated with poly(A) polymerase (PAP). Gene specific primer (GSP) 1 and 2 were used in PCR1 and nested PCR, respectively. In this experiment, the mamRNA short isoforms ended one nucleotide shorter than currently annotated, with a G that could correspond either to the first nucleotide of the intron, or to the first nucleotide of exon2, in which case the 3’-end of the isoform matched the predicted 5’ Pac1 cleavage site.

**Supplementary Figure 4:**
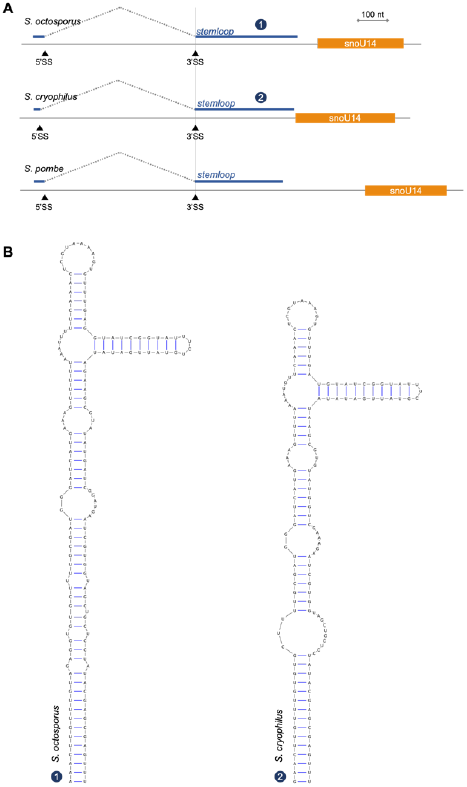
**A**. Genomic arrangements of snoU14 orthologs in *Schizosaccharomyces* species, aligned to their upstream 3’splice-site (3’SS) **B**. Predicted secondary structure of the sequences spanning the exon-exon junction in (A) except for S. pombe snoU14, already shown in main Figure 2.A.

## References

1. Dupuis-Sandoval, F., Poirier, M. & Scott, M. S. The emerging landscape of small nucleolar RNAs in cell biology. Wiley Interdiscip. Rev. RNA 6, 381 (2015).

2. Huang, Z. hao, Du Y. ping, Wen J. tao, Lu B. feng & Zhao, Y. snoRNAs: functions and mechanisms in biological processes, and roles in tumor pathophysiology. Cell Death Discov. 2022 81 8, 1– 10 (2022).

3. Gallie, D. R. The cap and poly(A) tail function synergistically to regulate mRNA translational efficiency. Genes Dev. 5, 2108–2116 (1991).

4. Grzechnik, P. et al. Nuclear fate of yeast snoRNA is determined by co-transcriptional Rnt1 cleavage. Nat. Commun. 9, (2018).

5. Kufel, J. & Grzechnik, P. Small Nucleolar RNAs Tell a Different Tale. Trends Genet. 35, 104–117 (2019).

6. Fafard-Couture, É., Labialle, S. & Scott, M. S. The regulatory roles of small nucleolar RNAs within their host locus. RNA Biol. 21, 1– 11 (2024).

7. Bergeron, D. et al. snoDB 2.0: an enhanced interactive database, specializing in human snoRNAs. Nucleic Acids Res. 51, D291– D296 (2023).

8. Larochelle, M. et al. Common mechanism of transcription termination at coding and noncoding RNA genes in fission yeast. Nat. Commun. 9, (2018).

9. Lemay, J. F. et al. The Nuclear Poly(A)-Binding Protein Interacts with the Exosome to Promote Synthesis of Noncoding Small Nucleolar RNAs. Mol. Cell 37, 34–45 (2010).

10. Chanfreau, G., Legrain, P. & Jacquier, A. Yeast RNase III as a key processing enzyme in small nucleolar RNAs metabolism. J. Mol. Biol. 284, 975–988 (1998).

11. Lee, C. Y., Lee, A. & Chanfreau, G. The roles of endonucleolytic cleavage and exonucleolytic digestion in the 5’-end processing of S. cerevisiae box C/D snoRNAs. RNA 9, 1362–1370 (2003).

12. Terns, M. P. & Dahlberg, J. E. Retention and 5′ Cap Trimethylation of U3 snRNA in the Nucleus. Science (80-. ). 264, 959–961 (1994).

13. Ghazal, G., Ge, D., Gervais-Bird, J., Gagnon, J. & Abou Elela, S. Genome-Wide Prediction and Analysis of Yeast RNase III-Dependent snoRNA Processing Signals. Mol. Cell. Biol. 25, 2981– 2994 (2005).

14. Yague-Sanz, C., Duval, M., Larochelle, M. & Bachand, F. Cotranscriptional RNA cleavage by Drosha homolog Pac1 triggers transcription termination in fission yeast. Nucleic Acids Res. 49, 8610–8624 (2021).

15. Rotondo, G., Huang, J. & Frendewey, D. Substrate structure requirements of the Pac1 ribonuclease from Schizosaccharomyces pombe. RNA 3, 1182–1193 (1997).

16. Ivakine, E., Spasov, K., Frendewey, D. & Nazar, R. N. Functional significance of intermediate cleavages in the 3′ETS of the pre-rRNA from Schizosaccharomyces pombe. Nucleic Acids Res. 31, 7110–7116 (2003).

17. Lamontagne, B. & Elela, S. A. Evaluation of the RNA Determinants for Bacterial and Yeast RNase III Binding and Cleavage. J. Biol. Chem. 279, 2231–2241 (2004).

18. Andric, V. et al. A scaffold lncRNA shapes the mitosis to meiosis switch. Nat. Commun. 12, (2021).

19. Leroy, E. et al. A bifunctional snoRNA with separable activities in guiding rRNA 2’-O-methylation and scaffolding gametogenesis effectors. bioRxiv 2024.10.03.615557 (2024). doi:10.1101/2024.10.03.615557

20. Wood, V., Lock, A., Rutherford, K. & Harris, M. PomBase lzlzlz the scientific resource for fission yeast. 1757, (2017).

21. Thodberg, M. et al. Comprehensive profiling of the fission yeast transcription start site activity during stress and media response. Nucleic Acids Res. 47, 1671–1691 (2019).

22. Zhang, Y. & Pelechano, V. High-throughput 5′P sequencing enables the study of degradation-associated ribosome stalls. Cell Reports Methods 1, 100001 (2021).

23. Samarsky, D. A., Schneider, G. S. & Fournier, M. J. An essential domain in Saccharomyces cerevisiae U14 snoRNA is absent in vertebrates , but conserved in other yeasts. 24, 2059–2066 (1996).

24. Zagorski, J., Tollervey, D. & Fournier, M. J. Characterization of an SNR gene locus in Saccharomyces cerevisiae that specifies both dispensible and essential small nuclear RNAs. Mol. Cell. Biol. 8, 3282–3290 (1988).

25. Harrison, P. W. et al. Ensembl 2024. Nucleic Acids Res. 52, D891– D899 (2024).

26. Chanfreau, G., Rotondo, G., Legrain, P. & Jacquier, A. Processing of a dicistronic small nucleolar RNA precursor by the RNA endonuclease Rnt1. EMBO J. 17, 3726–3737 (1998).

27. Yin, Q. F. et al. Long noncoding RNAs with snoRNA ends. Mol. Cell 48, 219–230 (2012).

28. Liu, Y. et al. Splicing inactivation generates hybrid mRNA-snoRNA transcripts targeted by cytoplasmic RNA decay. Proc. Natl. Acad. Sci. U. S. A. 119, e2202473119 (2022).

29. Gagnon, J., Lavoie, M., Catala, M., Malenfant, F. & Elela, S. A. Transcriptome Wide Annotation of Eukaryotic RNase III Reactivity and Degradation Signals. PLoS Genet. 11, 1–29 (2015).

30. Danin-Kreiselman, M., Lee, C. Y. & Chanfreau, G. RNAse III-mediated degradation of unspliced pre-mRNAs and lariat introns. Mol. Cell 11, 1279–1289 (2003).

31. Mattioli, C., Pianigiani, G. & Pagani, F. A competitive regulatory mechanism discriminates between juxtaposed splice sites and pri-miRNA structures. Nucleic Acids Res. 41, 8680–8691 (2013).

32. Melamed, Z. et al. Alternative splicing regulates biogenesis of miRNAs located across exon-intron junctions. Mol. Cell 50, 869– 881 (2013).

33. Havens, M. A., Reich, A. A. & Hastings, M. L. Drosha Promotes Splicing of a Pre-microRNA-like Alternative Exon. PLoS Genet. 10, 1004312 (2014).

34. Lee, D., Nam, J. W. & Shin, C. DROSHA targets its own transcript to modulate alternative splicing. RNA 23, 1035–1047 (2017).

35. Kim, B., Jeong, K. & Kim, V. N. Genome-wide Mapping of DROSHA Cleavage Sites on Primary MicroRNAs and Noncanonical Substrates. Mol. Cell 66, 258-269.e5 (2017).

36. Zhang, X. R., He, J. B., Wang, Y. Z. & Du, L. L. A cloning-free method for CRISPR/Cas9-mediated genome editing in fission yeast. G3 Genes, Genomes, Genet. 8, 2067–2077 (2018).

37. Qiu, C. et al. Thiolutin has complex effects in vivo but is a direct inhibitor of RNA polymerase II in vitro. Nucleic Acids Res. 52, 2546–2564 (2024).

38. Grigull, J., Mnaimneh, S., Pootoolal, J., Robinson, M. D. & Hughes, T. R. Genome-Wide Analysis of mRNA Stability Using Transcription Inhibitors and Microarrays Reveals Posttranscriptional Control of Ribosome Biogenesis Factors. Mol. Cell. Biol. 24, 5534 (2004).

39. Shomron, N., Malca, H., Vig, I. & Ast, G. Reversible inhibition of the second step of splicing suggests a possible role of zinc in the second step of splicing. Nucleic Acids Res. 30, 4127 (2002).

40. Rotondo, G. & Frendewey, D. Purification and characterization of the Pac1 ribonuclease of Schizosaccharomyces pombe. Nucleic Acids Res. 24, 2377–2386 (1996).

41. Horowitz, B., Goldfinger, B. A. & Marmur, J. Effect of cordycepin triphosphate on the nuclear DNA-dependent RNA polymerases and poly(A) polymerase from the yeast, Saccharomyces cerevisiae. Arch. Biochem. Biophys. 172, 143–148 (1976).

42. Bähler, J. & Wise, J. A. Preparation of Total RNA from Fission Yeast. Cold Spring Harb. Protoc. 2017, pdb.prot091629 (2017).

43. Yague-Sanz, C. et al. Chromatin remodeling by Pol II primes efficient Pol III transcription. Nat. Commun. 14, (2023).

44. Liu, Q., Li, X. & Sommer, S. S. pK-matched running buffers for gel electrophoresis. Anal. Biochem. 270, 112–122 (1999).

45. Duval, M. et al. The conserved RNA-binding protein Seb1 promotes cotranscriptional ribosomal RNA processing by controlling RNA polymerase I progression. Nat. Commun. 14, (2023).

46. Hofacker, I. L. Vienna RNA secondary structure server. Nucleic Acids Res. 31, 3429–3431 (2003).

47. Darty, K., Denise, A. & Ponty, Y. VARNA: Interactive drawing and editing of the RNA secondary structure. Bioinformatics 25, 1974– 1975 (2009).

48. Li, H. Minimap2: pairwise alignment for nucleotide sequences. Bioinformatics 34, 3094–3100 (2018).

49. Thorvaldsdottir, H., Robinson, J. T. & Mesirov, J. P. Integrative Genomics Viewer (IGV): high-performance genomics data visualization and exploration. Br. Bioinform 14, 178–192 (2013).

